# Morphological and functional characterization of the ptychocyte, a stingless stinging cell

**DOI:** 10.64898/2026.04.10.717713

**Authors:** Ana Hoffman Sole, Kennedy Bolstad, Elijah James, Chris Roh, Leslie S. Babonis

## Abstract

Cnidocytes (stinging cells), unique to cnidarians (corals, anemones, jellyfish), have diversified into distinct types with variable forms and functions. Nematocytes, cnidocytes found in all cnidarians, are used for prey capture and defense. When triggered, a pressurized capsule inside the nematocyte releases a harpoon-like structure attached to a hollow tubule that pierces prey and delivers venom. Ptychocytes, a cnidocyte unique to tube anemones (sister to corals and sea anemones) discharge a long spineless tubule used exclusively to build the tube in which the animal lives. Given that nematocytes and ptychocytes are specialized for different functions, we hypothesized that they might respond to firing cues in different ways. To test this, we examined the morphology, function, and distribution of nematocytes and ptychocytes in the North American Tube Anemone, *Ceriantheopsis americana*. We determined that ptychocytes have apical sensory structures like the cones previously described on nematocytes. Surprisingly, the body wall has a dense population of multiciliated cells that appear to function in tube formation. To determine how divergent selection pressures may have affected firing dynamics, we compared the discharge kinematics of cnidocytes from *C. americana* and the model sea anemone, *Nematostella vectensis*. Both nematocytes and ptychocytes from *C. americana* fired slower than nematocytes from *N. vectensis*, suggesting the rapid discharge speed of sea anemone nematocytes resulted from modification to these cells after sea anemones and tube anemones diverged from their common ancestor. By comparing the morphology and function of different cnidocytes, we can reconstruct the steps that gave rise to cnidocyte diversity.

## Introduction

Cnidocytes (stinging cells) are a diverse group of cells unique to cnidarians (the clade including sea anemones, corals, and jellyfish). Highly variable in form and function, cnidocytes are an important model for cell type diversification (Babonis 2025; Babonis and Martindale 2014). All cnidocytes have a stinging organelle, composed of a pressurized capsule containing an eversible tubule. When the cnidocyte fires, the capsule opens and the release of pressure causes the tubule to evert (Tardent 1995). Since their origin in the stem cnidarian ∼700-800 million years ago (McFadden et al. 2021; Sierra and Gold 2024), cnidocytes have diversified into three broad types: nematocytes, spirocytes, and ptychocytes (Fig 1A). In nematocytes, which are found in all cnidarians, the capsule has a thick wall, and the eversible tubule has been modified (in many types of nematocytes) to act as a harpoon, delivering venom to immobilize prey and deter predators (Mariscal 1984). Spirocytes, which are restricted to hexacorals (hard corals, sea anemones, tube anemones, and their relatives), have a thin capsule wall surrounding a long, helically coiled, tubule used to ensnare prey (Mariscal et al. 1976). Ptychocytes, which are unique to tube anemones, have a thin capsule wall and a tubule packed in a pleated fold like a firehose (Mariscal et al. 1977). Unlike nematocytes and spirocytes, which are used primarily to capture prey, ptychocytes are used exclusively to construct the tube in which tube anemones live. Tube anemones are, therefore, the only cnidarian with all three types of cnidocytes, which makes this clade of animals particularly useful for understanding the diversification of spirocytes and ptychocytes from a putative nematocyte-like ancestral cnidocyte.

**Figure 1.**
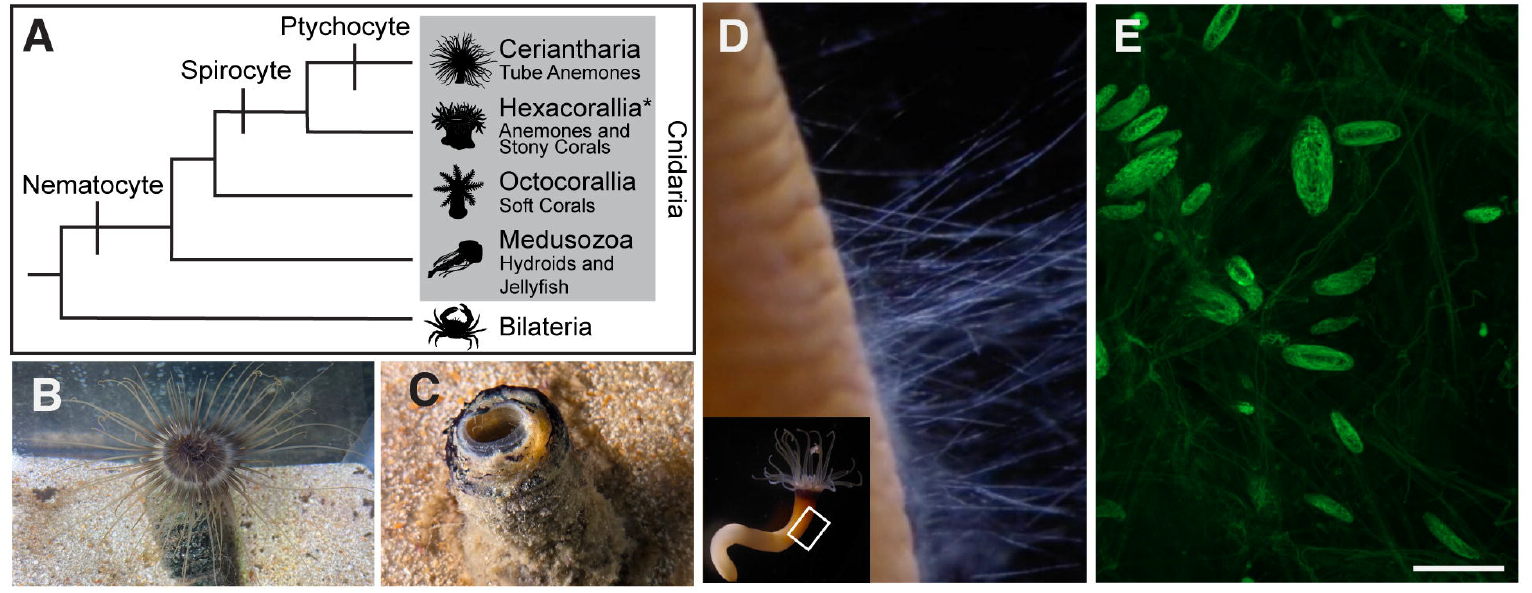
Ptychocytes are unique to tube anemones. **A)** Cladogram of cnidarians showing hypothesized origin of each cnidocyte type. The branch labeled Hexacorallia* is currently unnamed and represents all non-Ceriantharian hexacorals. **B)** *C. americana* with the tentacles and oral disc projecting out from the tube. **C)** *C. americana* retracted into the tube, tentacles and oral disk are no longer visible. **D)** High magnification image extracted from a video of ptychocytes firing from the body wall of *C. americana* after it was removed from its tube (inset shows whole animal with tube removed). **E)** Ptychocytes are autofluorescent and can be easily seen in the body wall of an animal removed from its tube. Confocal z-stack maximum projection, scale bar is 50um.

Tube anemones are sister to the rest of hexacorals (the group containing stony corals, sea anemones, and their relatives) and have received less scientific focus than these taxa. There are approximately 50 accepted species of tube anemones found throughout the world’s oceans where they live partially buried into soft sediment in the sea floor (Stampar et al. 2020; Forero Mejia et al. 2020). These animals are named for the membranous tubes that they make and withdraw into when disturbed (Fig 1B,C). The tube is primary composed of fired ptychocytes (Fig 1D,E) but can also incorporate fired nematocytes and spirocytes, as well as mucus and sediment from the animal’s environment (Frey 1970; Mariscal et al. 1977; Stampar, Beneti, et al. 2015). Tube anemones continue to make tubes throughout their lifetime to accommodate growth, and an adult removed from its tube will begin firing ptychocytes to build a new tube within minutes of removal (Mariscal et al. 1977; Stampar et al. 2015, Fig 1D). Tube anemones have two types of tentacles used to capture prey: an outer ring of long marginal tentacles populated by nematocytes and spirocytes, and an inner ring of short labial tentacles both populated by abundant nematocytes (microbasic b-mastigophores) and spirocytes (Widersten 1976; Forero-Mejia et al. 2024). Internally, tube anemones have gonad-forming mesenteries extending aborally from the pharynx and thread-like mesenteries (sometimes referred to as craspedonemes in other tube anemones; McMurrich 1910; Forero-Mejia et al. 2024) that extend into the body cavity from the oral end of the gonad-forming mesenteries. While the nematocytes from sea anemones are known to use mechanosensory and chemosensory information as a firing cue, the cue used to trigger cnidocytes in tube anemones has not been investigated.

Sea anemones have served as important models for understanding the physiological control of cnidocyte discharge (Watson et al. 1998). Previous studies show that sea anemone nematocytes have apical sensory structures that help them respond rapidly to vibrational and chemical cues from their environment (Watson and Hessinger 1989). These apical structures consist of a central cilium surrounded by a cone of microvilli. Extensive work on the nematocytes of sea anemones has revealed the importance of these apical sensory cones for tuning nematocyte firing to a frequency that closely matches the movement pattern of passing prey (Watson and Hessinger 1989). Spirocytes, notably, have no apical sensory cone (Mariscal and Bigger 1976). In sea anemones, spirocytes are connected to neighboring nematocytes by a synapse, and it is thought that this helps trigger spirocytes to fire synchronously (Weir et al. 2020). It seems possible that because ptychocytes are not used to capture prey, they might also lack an apical sensory cone (like spirocytes) and may not be innervated by nearby nematocytes. Since they were first described (Mariscal et al. 1977), there have been very few studies of ptychocytes and these studies have mostly focused on their morphology, distribution, and value as a taxonomic character (Forero-Mejia et al. 2024; Reft and Daly 2012; Stampar et al. 2016; Fautin 2009). There has been limited investigation of ptychocytes in situ so little is known about the structures that support them. Likewise, there are no reports of the firing mechanics of live ptychocytes, resulting in a gap in our knowledge of the relationship between structure and function across cnidocytes.

To better characterize the functional diversity of cnidocytes, we investigated the morphology, in situ environment, and firing dynamics of ptychocytes from the North American Tube Anemone, *Ceriantheopsis americana*. While some tube anemones have long planktonic larval stages (Stampar, Morandini, et al. 2015), *C. americana* does not, so it is easily maintained in the lab. We show that *C. americana* has a diverse array of cnidocytes and that ptychocytes are restricted to the body wall of this animal. Furthermore, ptychocytes have apical sensory cones, similar to those found on nematocytes, and speed of tubule discharge is not significantly different between nematocytes and ptychocytes from *C. americana*, but both are significantly slower than nematocytes from the sea anemone *Nematostella vectensis*.

## Methods

### Care and culture of the North American Tube Anemone

Tube anemones were collected with their tubes at low tide from the public beach in Cedar Key, FL (29.13544, -83.03701). Animals were maintained in artificial seawater (27-30ppt, Tropic Marin Pro Reef Salt, Wartenberg, Germany) at room temperature (22°C) with a 12/12 light/dark cycle and provided with about six inches of sand to support their tubes. Animals were fed weekly with chopped black tiger shrimp (*Penaeus monodon*) and supplemented with freshly hatched *Artemia salina* twice per week.

### Tissue collection and quantitative cnidocyte analysis

Like other cnidarians, the North American tube anemone appears to be highly regenerative. To collect tissue samples, animals were removed from their tubes, dissected using sharp forceps and a microscalpel, then returned to their tank to regenerate. Dissected tissues were fixed for 90 seconds in 4% paraformaldehyde with 0.2% glutaraldehyde in phosphate-buffered saline with 0.1% Tween-20 (PTw) at room temperature (22°C). Initial fixative was removed before tissues were further fixed in 4% paraformaldehyde in PTw (no glutaraldehyde) for one additional hour at 4°C. To remove fixative, tissues were washed three times in PTw and stored tissues in clean PTw at 4°C before processing.

To quantify the types and proportions of cnidocytes across different tissues, each piece of tissue was placed on a glass slide with a drop of 80% glycerol (diluted in PTw). To spread cells out across the slide, a coverslip was pressed down on the tissue and gently rubbed across the slide. Nail polish was applied to the edge of the coverslips so that slides could be stored and imaged indefinitely. Cnidocytes were quantified along three or more non-overlapping visual transects of each slide viewed on a compound microscope (Nikon Eclipse E800). We analyzed tissues from three individuals, collecting three samples of each tissue from each individual from the tip, middle, and base of the marginal tentacles, the tip and middle of the labial tentacles, thread-like mesentery, gamete forming mesentery, and the body wall.

### Immunohistochemistry

To characterize the apical sensory structures found on cnidocytes, we used immunohistochemistry to fluorescently label proteins. Tissue was collected and fixed as for quantitative cnidocyte analysis before being incubated for one hour in phosphate buffered saline with 0.2% Triton X-100 and 0.1% BSA (PBT). Tissues were removed from this solution and incubated in 5% normal goat serum (diluted in PBT) for 1 hour at room temperature (22°C), then incubated at 4°C for at least 20 hours in primary antibody diluted in 5% normal goat serum in PBT. Cilia were labeled using a commercially available mouse antibody recognizing acetylated tubulin (Sigma T6743) at 1:500 concentration. Primary antibody was removed with five washes in PBS with 0.2% Triton-X before tissues were incubated for 20 hours at 4°C in 1:500 goat anti mouse secondary antibody. To visualize F-actin in microvilli, we counterstained tissues with phalloidin (Molecular Probes #A12379) diluted 1/200 in PBS with 0.2% Triton-X. All tissues were mounted in 80% glycerol on slides coated with Rain-X (Illinois Tool Works, USA) and imaged on a Zeiss LSM 900 upright confocal microscope. Images were adjusted for brightness and contrast in Fiji (Schindelin et al. 2012).

### Scanning Electron Microscopy

To further characterize the apical morphology of cnidocytes and other sensory cells, tissues were imaged with scanning electron microscopy. Samples were fixed at least ten hours at 4°C in 2.5% Glutaraldehyde in 3X MarPHEM buffer (Montanaro et al. 2016) and then washed extensively in 3X MarPHEM buffer to remove the fixative. Tissues were washed in 0.05M cacodylate buffer three times for 10 minutes each before secondary fixation in 1% OsO_4_ for one hour at room temperature (22°C). To remove the secondary fixative, samples were washed three times for 10 minutes each in 0.05M cacodylate buffer and then samples were dehydrated by washing for 10 minutes each in a graded ethanol series: 25%, 50%, 70%, 95%, and then twice in 100%. To remove the rest of the moisture, tissues were critical point dried (Bal-Tec CPD 030) and then mounted on pin stub specimen mounts using double-stick carbon tape. Samples were sputter coated (3 minutes, 30mA, Au-Pd) in a Denton Desk V sputter coater (Denton Vacuum, Moorestown, NY, USA) and imaged with an environmental scanning electron microscope (JEOL JCM-7000 Neoscope).

### Live Cnidocyte Firing

To compare initial velocity of tubule eversion during firing in different cnidocyte types, cells were dissociated from live tissues, induced to fire, and captured with high-speed video. To dissociate cells, a small piece of live tissue was cut from the marginal tentacle (for *C. americana* nematocytes), body wall (for ptychocytes), or the tentacles (for *N. vectensis* nematocytes) and washed twice in calcium/magnesium-free artificial sea water to remove excess divalent ions. Tissues were then incubated in clean calcium/magnesium-free seawater for at least one hour at room temperature and subjected to gentle agitation (flicking) approximately every ten minutes. To induce firing in isolated cnidocytes, a drop of dissociated cells was pipetted into the bottom of a 35mm glass-bottom petri dish (VWR #10810-054) and compressed under a coverslip supported on clay “feet”. The dish was filled with 1mL of artificial sea water and mounted on the stage of an inverted compound microscope (Nikon, Inverted eclipse Ti) fitted with a high-speed camera (Phantom, Vision Research Phantom v711). Firing was induced by adding acetic acid to the seawater to a final concentration of 0.15% and videos were captured at 2000 frames per second.

### Video Analysis

Cnidocyte kinematics were generated by first tracking tip position in Tracker Video Analysis and Modeling Tool v6.3.3 then calculating kinematics in MATLAB R2024b. In Tracker, pre-recorded cnidocyte firing events were imported and point masses were assigned to the tip as well as the capsule at each frame of the firing event. Then the raw pixel coordinates in the video frame (labeled *pixelx* and *pixely* in Tracker when extracting) and the frame numbers were extracted to be imported into MATLAB.

In MATLAB, the raw pixel coordinates were imported to calculate relative kinematics. First, all pixel coordinates of the capsule were subtracted from that of the tip to ensure that any kinematics calculated are that of the tip relative to the capsule. Then, the raw pixel coordinates were converted to physical units (i.e. micrometers) using the microscope magnification and frame indices were converted to time in seconds using the recording frame rate. The x- and y-components of relative velocity and acceleration were calculated using central differencing:

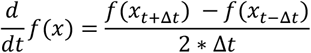

When calculating velocity, *f(x)* refers to the tip position while *Δt = t*_*2*_*-t*_*1*_. Meanwhile, when calculating acceleration *f(x)* refers to the tip velocity. The resultant vector magnitudes of the kinematics were calculated from their respective components:

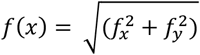

Statistical analysis was conducted for the average velocity of each cell type for the first 0.05 seconds in MATLAB. A one-way ANOVA was computed for an alpha of 0.05 with the *anova1* function. Then, the Tukey-Kramer test was performed using the *multcompare* function to compare between all possible pairs.

## Results

### *C. americana* has a diverse suite of cnidocytes

We found 15 types of cnidocytes across all the examined tissues (Fig 2A-C): Seven were nematocytes (Fig 2D-J), four were gracile spirocytes (Fig 2K-N), and four were ptychocytes (Fig 2O-R). The descriptions provided here follow the nomenclature of Östman et al. (2010) and Forero-Mejia et al. (2024). We found two types of holotrichous isorhizas, a large one with a thick tubule, clearly visible through the capsule (Fig 2D), and a small one with a thin tubule, barely visible through the capsule (Fig 2E). We also found three types of microbasic b-mastigophores: a small one with a harpoon more than half the length of the capsule (Fig 2F), a large one with a harpoon more than half the length of the capsule (Fig 2G), and one with the harpoon taking up about half the capsule (Fig 2H). We found two types of basitrichous isohrizas: one with a long harpoon (Fig 2I) and one short (Fig 2J). Four of the cnidocyte types were gracile spirocytes, which we distinguished by two features: size and the presence of an empty space between the tubule and capsule on the end of the organelle opposite the apex (Fig 2K-N) We identified four types of ptychocytes, which we could reliably distinguish by the presence of an apical bend and their approximate length to width ratio (Fig 2O-R).

**Figure 2.**
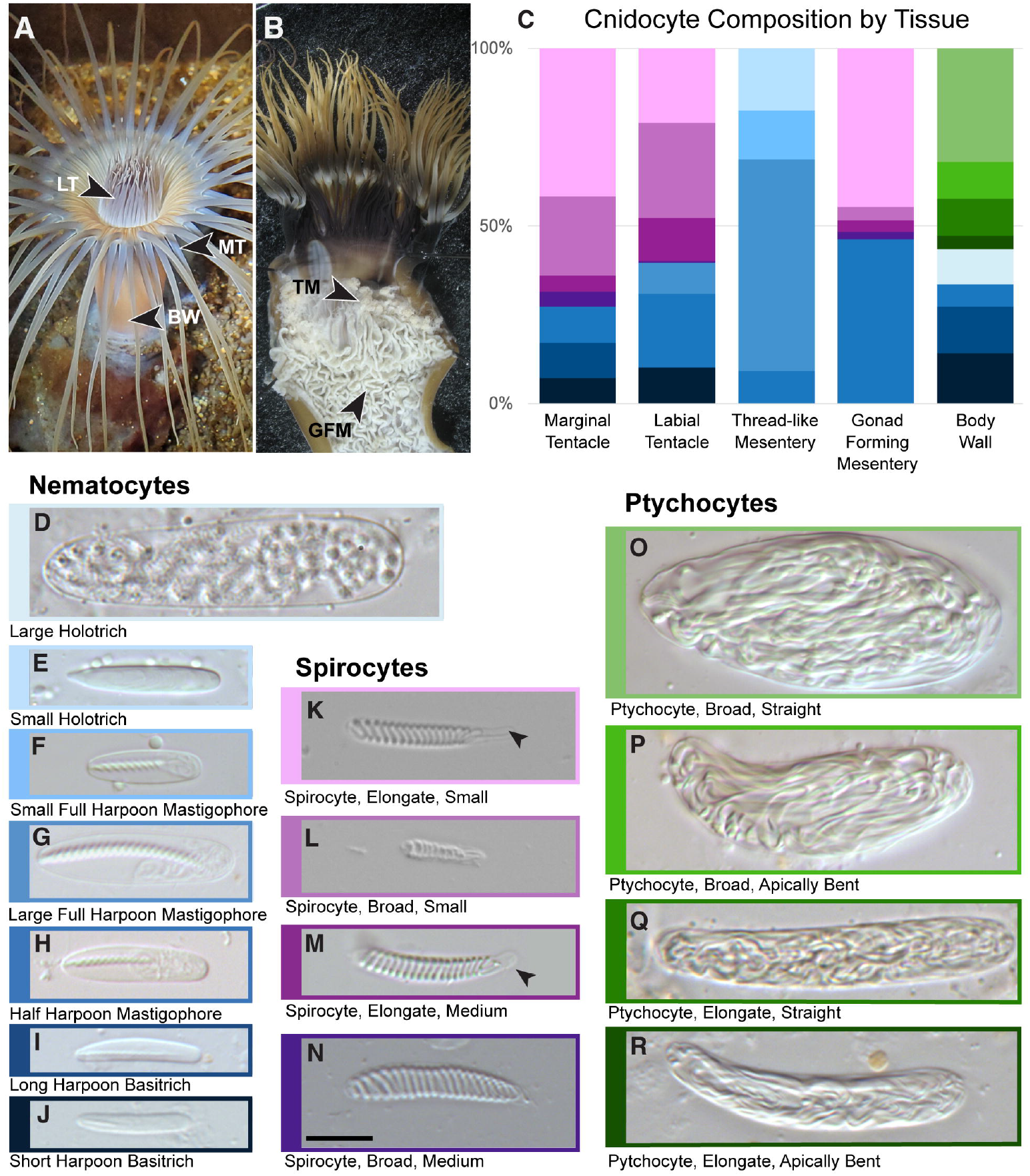
Diversity and distribution of cnidocytes in *C. americana*. **A,B)** Live animal showing locations of examined tissues. BW – body wall, TM – thread-like mesentery, GFM – gonad forming mesentery, LT – labial tentacle, MT – marginal tentacle. **C)** Cnidocyte composition by tissue, colors correspond to outlines of images D-R. N = 3 individuals per tissue. **D-R)** Differential interference contrast (DIC) images of isolated cnidocytes. **K,M)** Arrows point to space in elongate spirocyte capsules opposite end from which eversion occurs. Scale bar = 10um and applies to all DIC images.

### Ptychocytes are restricted to the body wall

As previously described (Widersten 1976), we found both the marginal and labial tentacles (Fig 2A) to be populated by spirocytes and nematocytes. The marginal tentacles have both kinds of basitrichous isohrizas, half harpoon microbasic b-mastigophores, and all four types of spirocytes (Fig 2C). Nematocytes compose only 28% of the cnidocytes present, so the tissue is dominated by spirocytes. Of the spirocytes, small elongate spirocytes were the most common, followed by small broad spirocytes. Labial tentacles had a similar composition (40% nematocytes) but had a different suite of nematocytes: the short basitrichous isohriza, half harpoon microbasic b-mastigophore, and full harpoon microbasic b-mastigophore. The thread-like mesentery, found inside the body on near the pharynx, contains only nematocytes: three types of mastigophores and the small holotrichous isohriza, which is restricted to this tissue. The full harpoon microbasic b-mastigophore was the most abundant cell in this tissue. The gonad forming mesentery is found along most of the inside of the animal, starting below the thread-like mesentery (Fig 2B). Remarkably, it contained all four types of spirocytes as well as the half harpoon microbasic b-mastigophore.

The body wall is dominated by ptychocytes (Fig 2C). Ptychocytes make up 56% of the cnidocytes in the body wall, with broad cells being more common than narrow ones. The other half (44%) of the cnidocytes found in the body wall were nematocytes, and included large holotrichous isohrizas which, like ptychocytes, were also exclusively found in this tissue. The four types of ptychocytes and the large holotrichous isohrizas are the five largest cnidocytes present in *C. americana*, and are all restricted to the body wall. The rest of the nematocytes present in the body wall were also found in other tissues.

### Ptychocytes have apical sensory cones similar to nematocytes

To learn more about the mechanisms that control cnidocyte firing, we examined the apical sensory structures on different types of cnidocytes and across different tissues in *C. americana*. Initial scanning electron microscope images showed that the body wall is covered in cilia that obscure other features on the surface of the tissue (Fig 3A). Further examination of these cilia with immunofluorescence revealed that they emerge from multiciliated cells that dominate the outer surface of the body wall (Fig 3B,C). Each multiciliated cell has a ring of short actin-based microvilli marking the boundary of the cell, which makes it possible to see that 9 to 12 cilia protrude from each cell.

**Figure 3.**
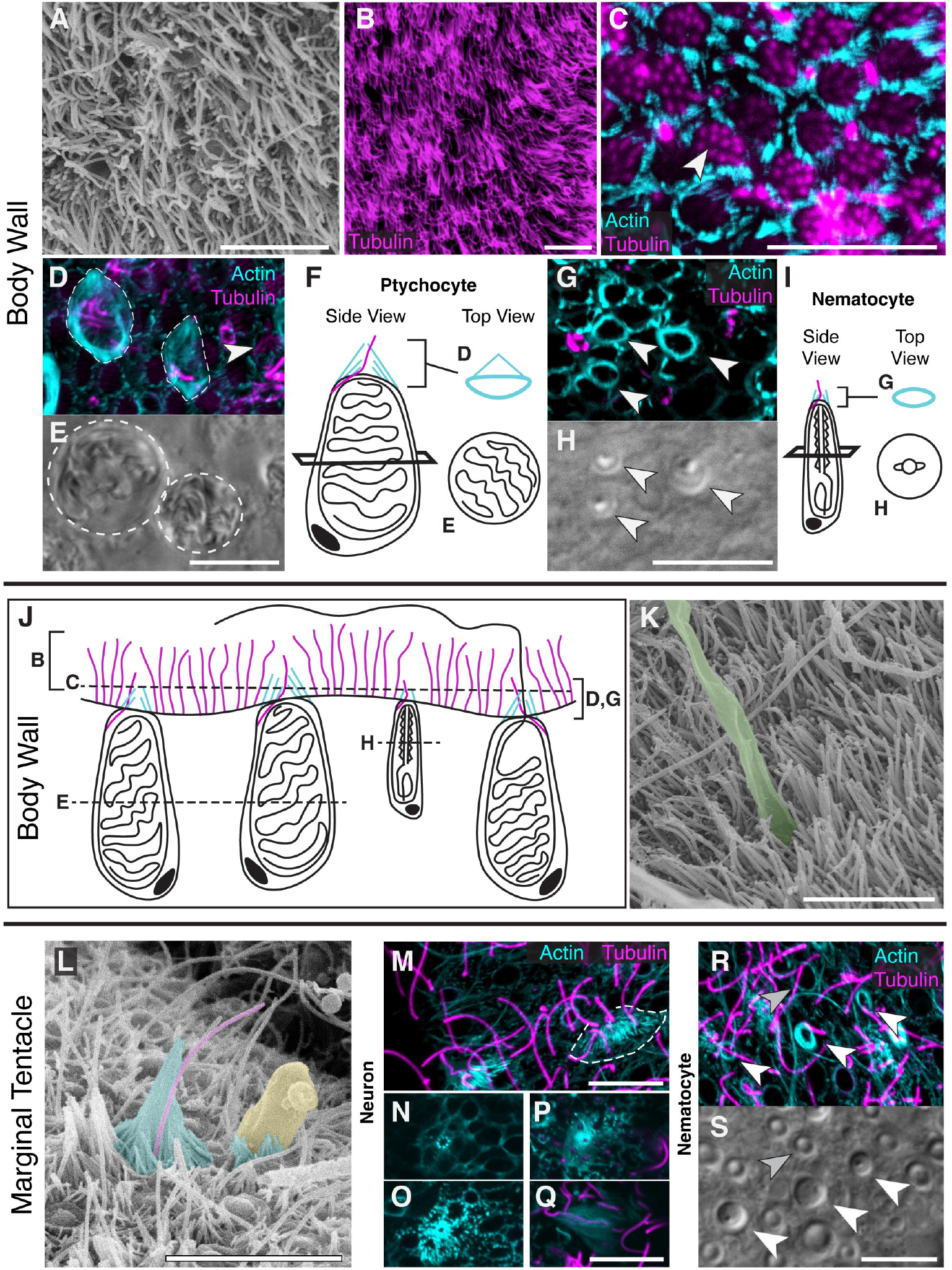
Examination of cnidocytes in situ in the body wall and marginal tentacle. **A,B,C)** Multiciliated cells in the body wall. **A,B)** Abundant cilia on the surface of the body wall examined by SEM and labeled with anti-acetylated tubulin antibody (magenta). **C)** A deeper focal plane of the region shown in B showing a cluster of multiciliated cells each with approximately 11 cilia emerging from the apex (white arrowhead). Actin in the cell boundaries is labeled by phalloidin (cyan). **D,E)** Deeper focal planes through a region of body wall showing two microvilli cones (outlined in dashed line) on top of ptychocytes. An adjacent multiciliated cell is indicated by the white arrowhead. (E) Ptychocytes, are recognizable by their capsule morphology shown here using DIC. **F)** Schematic representation of the structures shown in D,E. **G)** Similar cones on top of nematocytes (white arrowheads), which are recognizable by their capsule morphology (H). **I)** Schematic summarizing the structures shown in G,H. **J)** Schematic of body wall tissue. **K)** SEM of fired ptychocyte tubule (false-colored green) emerging from underneath a field of cilia in body wall. **L)** SEM of marginal tentacle with tall, sensory neuron cone (left, false-colored cyan) with a cilium (magenta) next to a nematocyte capsule (yellow) being squeezed out of the tissue through a short microvilli cone (cyan). **M)** Tall cones in marginal tentacle with a central cilium (confocal max projection). **N-Q)** A series of optical sections through a tall cone, showing multiple cones with cilia contributing to composite cone, lettered from deepest in the tissue to surface. **R,S)** Short microvilli cones (cyan) in marginal tentacles on top of nematocytes (white arrowheads), but not on top of spirocytes (gray arrowhead). Scale bars all 10um.

Using optical sectioning of fluorescently labeled tissues and associated DIC images, we looked at the structures on top of the cnidocytes adjacent to these multiciliated cells. We observed cones of microvilli surrounding a central cilium on top of both ptychocytes and nematocytes (Fig 3D-I). Using associated DIC slices, we confirmed that all microvillar cones observed in the body wall were associated with a cnidocyte, identifiable by the morphology of its stinging apparatus. Because of the abundance and length of the cilia emerging from the multiciliated cells, in SEM the position of the ptychocytes could only be determined when we saw discharged tubules extending out beyond the mat of cilia (Fig 3J,K).

We examined marginal tentacles to compare nematocyte and spirocyte apical sensory structures. Initial scanning electron microscope images showed many cones on the surface of the tenacle but did not indicate what cells were underneath these cones. The largest cones on the marginal tentacles were composite cones comprised of a central cilium supported by cones of microvilli from multiple cells (Fig 3L-Q). This cone morphology was previously reported to be on top of mechanosensory neurons in tube anemones (Peteya 1975) and other anthozoans (Westfall et al. 1998). One key difference between mechanosensory neuron cones described previously and the cones reported here is that here we see more than just the central kinocilium. Each cone contributed to the composite cone has a cilium, so cilia emerge from multiple places in the composite cone (Fig 3P). Nematocytes in the marginal tentacle had cones consistent with those seen on body wall nematocytes and spirocytes, as expected, had no cones at all (Fig 3R,S).

### Both ptychocytes and nematocytes fire slowly in *C. americana*

Given that the apical sensory morphology of ptychocytes was similar to what was observed for nematocytes, we explored whether these two cell types exhibited similar firing kinematics. Nematocyte discharge is known to be one of the fastest movements in biology (Nüchter et al. 2006). To determine if ptychocytes are just as fast as nematocytes, we examined the initial velocity of cnidocyte firing in ptychocytes and nematocytes (long harpoon basitrichous isohrizas) from *C. americana* and compared both to nematocytes (tentacle basitrichous isohrizas) from the sea anemone *Nematostella vectensis*(Fig 4A-D). In the first 0.05 seconds of firing, *N. vectensis* basitrichous isorhizas rapidly everted about one capsule length of tubule and then began to slow down (Fig 4E, F black curves), while both *C. americana* cnidocytes moved at about the same speed the whole time as evident from the red and blue curves Fig 4F. Over this period, ptychocytes and nematocytes from *C. americana* were both significantly slower than nematocytes from *N. vectensis* (Fig 4F,G). We found no significant difference between ptychocytes and nematocytes from *C. americana* (p = 0.23), but we did find that ptychocytes were significantly slower than N. vectensis nematocytes (p = 1.76 × 10^-6^) and *C. americana* nematocytes were also slower than *N. vectensis* nematocytes (p = 2.88 × 10^-5^).

**Figure 4.**
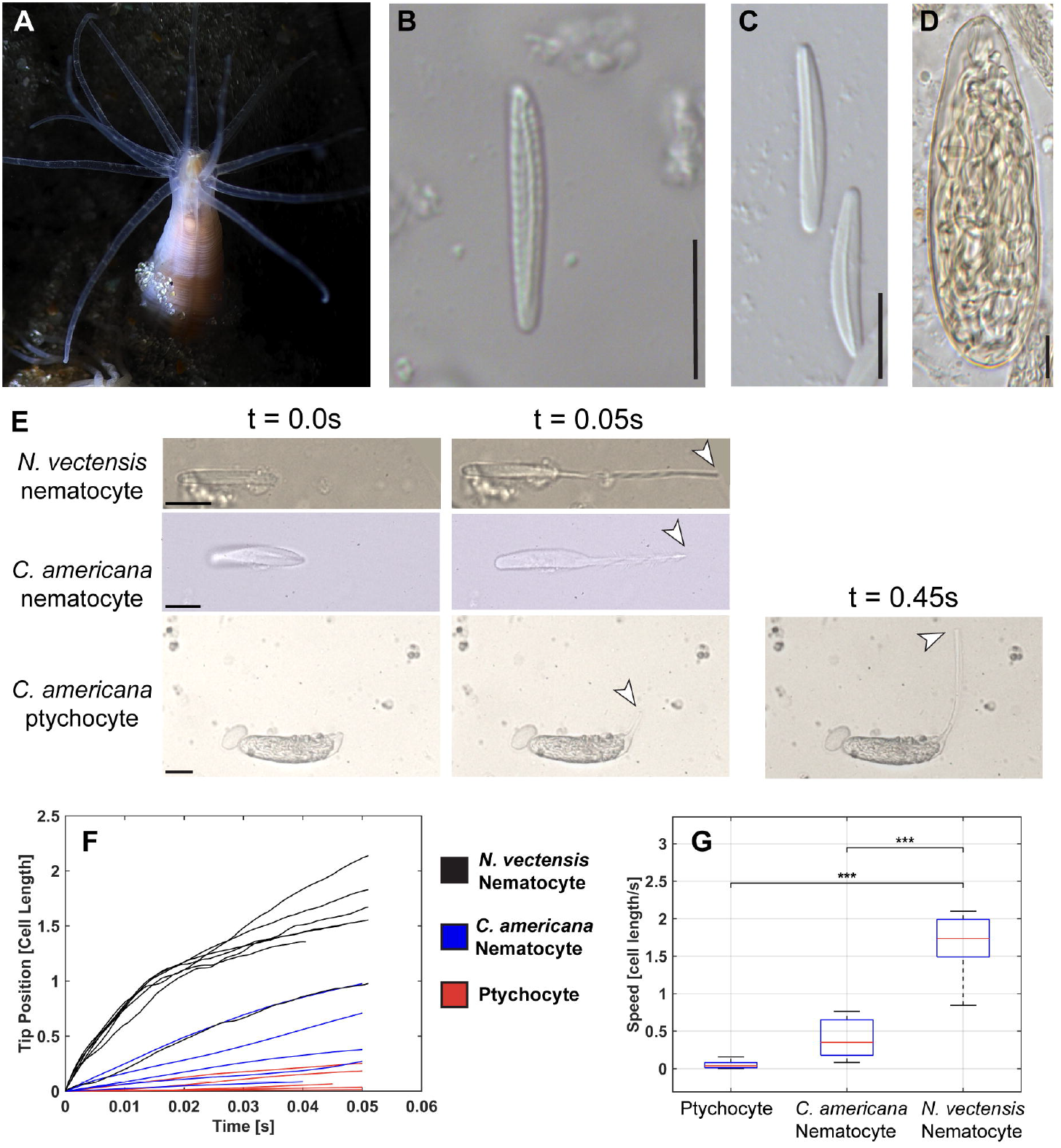
Tube anemone cnidocytes fire slowly **A)** The sea anemone *Nematostella vectensis* pictured burrowing in sand. **B)** A large basitrich isolated from the tentacle of *N. vectensis*. **C)** Two long harpoon basitrichs isolated from the marginal tentacle of *C. americana*. **D)** A broad, straight ptychocyte isolated from the body wall of *C. americana*. **E)** Still frames from high speed videos showing progression of tubule eversion. Images shown at start of firing where no eversion has happened yet, 0.05 seconds into firing (timeframe shown in F,G) where *N. vectensis* nematocyte has everted more than one capsule length and *C. american*a nematocyte has everted no more than one capsule length. Additional 0.45 second timepoint shows the ptychocyte, when it has everted one capsule length. **F)** Position of the tubule tip over the first 0.05 seconds of firing, normalized by cell length. N = 5 ptychocytes, N = 5 tube anemone nematocytes, N = 7 *N. vectensis* nematocytes. **G)** Box plot of average velocity in cell lengths per second over first 0.05 seconds of firing. Ptychocyte vs. *C. americana* nematocyte p = 0.23, Ptychocyte vs. *N. vectensis* nematocyte p = 1.76 × 10^-6^, *C. americana* nematocyte vs *N. vectensis* nematocyte p = 2.88 × 10^-5^. Scale bars all 10um.

## Discussion

### Cnidocyte regionalization enables functional specialization

Cnidarians have a seemingly simple body plan made of just two cell layers, an outer epidermis and an inner gastrodermis. However, within the morphological constraints of the diploblastic body plan, cnidarians achieve biological complexity through the diversification of cell types with unique functions. These unique cells are then restricted to relevant tissues, such that separate species may have similar body plans and morphologies, but different suites of cells used for different purposes. For example, many cnidarians have inducible organs of aggression used in intraspecific combat that are populated by a type of cnidocyte unique to those tissues (Williams 1991). In tube anemones, ptychocytes are restricted to the body wall, and perform a specific, and likely derived, function in constructing a tube (Fig 1,2). In ptychocytes, specialization for tube building is associated with a longer tubule, leading to single cells producing tubules upwards of two millimeters long (Mariscal et al. 1977). To accommodate such a long tubule, ptychocytes have larger capsules than most other cnidocytes and have tubules that fold down to enable better packing (Mariscal et al. 1977). As opposed to nematocytes which fire quickly to be able to pierce rapidly moving prey, it seems ptychocytes may not have evolved to maximize speed. Traits like long tubules, and thinner capsules might be tolerated in ptychocytes where they would not be in nematocytes. By investigating cell biology in the context of the behaviors of a species, we can learn more about cell type evolution and how novel cell functions might arise.

### Multiciliated cells are likely used to generate flow over the body wall

The body wall builds and maintains its own microenvironment, protected from the external environment by the surrounding tube. We describe for the first time a population of highly abundant motile multiciliated cells in the body wall of C. americana, (Fig 4). Multiciliated cells are present in representatives of many animal phyla including ctenophores, which are thought to be sister to the rest of metazoa (Dunn et al. 2008; Schultz et al. 2023). The morphology of multiciliated cells varies widely across taxa with some cells adorned by only 2-3 cilia whereas others contain hundreds (Purschke et al. 2017; Boutin and Kodjabachian 2019). Here, we show that *C. americana* has no more than 12 cilia per cell. This variation makes it difficult to determine if multiciliated cells were present in the common ancestor of all animals or if the ability to make cells with multiple cilia has arisen multiple times throughout the diversification of animals.

While we can observe that these multiciliated cells are motile, this precise purpose remains unresolved. Multiciliated cells are found across invertebrates and are most commonly used to create fluid flow for locomotion (Meunier and Azimzadeh 2016). The only other report of multiciliated cells from a cnidarian described cells with up 500 cilia per cell in a species of hydrozoan medusa, *Aglantha digitale*. These cells are found in bands along the tentacles and are used to increase water flow over the tentacles to improve prey capture (Mackie et al. 1989).

It is possible strong flow is advantageous to keep the space in between the body wall and the tube free from parasites. Alternatively, since multiciliated cells are only found in the same tissue as ptychocytes, it is possible that the flow created by these cells plays a role in tube construction. When nematocytes and spirocytes fire, they usually adhere to prey and the struggling prey helps pull the discharged stinging apparatus out of the cnidocyte membrane. Because ptychocytes do not land on prey, we suggest the adjacent multiciliated cells might create an equivalent source of flow pulling the discharged ptychocyte stinging apparatus out of the cell to facilitate formation of the mesh-like membrane of the tube. To test this hypothesis, cilia inhibitors could be applied to whole animals during tube construction, and the resulting tube could be compared with tube made by animals with uninhibited cilia.

### Body wall cnidocytes have different sensory support than tentacle cnidocytes

Tube anemone tentacle sensory structures seem to match our knowledge of sea anemone tentacle sensory structures. Both have mechanosensory neurons next to nematocytes and spirocytes, so we infer that tube anemones, like sea anemones, are using neurons to tune nematocyte firing to a frequency that matches passing prey. The body wall cnidocytes, however, are not used to catch prey in either taxon. We found no neuron sensory cones in the tube anemone body wall, and there are no reports in the literature of sensory neuron cones in the body wall of *N. vectensis* or any other sea anemone. This intriguing observation suggests that co-evolution of a cnidocyte and its neighboring cells may be important for diversifying the function of morphologically similar stinging cells found in different parts of the body, although this should be investigated further. Without sensory neuron cones present, it is possible tube anemone multiciliated cells may serve a sensory purpose. However, the apical sensory cones on the cnidocytes themselves may be sufficient when prey does not need to be caught. When a tube anemone is removed from its tube, it begins firing ptychocytes to construct a new one within minutes (Stampar, Beneti, et al. 2015; Mariscal et al. 1977). The different fluid dynamics experienced by multiciliated cells in open water (after the tube has been removed) rather than a confined space inside a fully formed tube could be the signal used to induce ptychocyte firing. Examination of the synapses between cnidocytes and surrounding cells in the body wall, similar to the work in Weir et al. (2020) could elucidate the relationships between these cells.

### Co-evolution of capsule structure and firing speed

Across the diversity of cnidocytes there is a unifying feature: each cell rapidly everts the content of the capsule when fired. Prior to the work presented here, investigation of the dynamics of cnidocyte discharge has been performed in only a small sample of cnidocyte types from a limited selection of taxa: Hydra (Nüchter et al. 2006; Holstein and Tardent 1984), two sea anemones (Karabulut et al. 2022; Godknecht and Tardent 1988) and a single species of jellyfish (Park et al. 2017). Nematocytes from medusozoans (hydroids and jellyfish) build up osmotic pressure in the capsule, causing a reinforced apical structure at the apex of the capsule (the operculum) to open and forcing the harpoon and tubule to evert, all of which is completed in under 0.003 seconds (Holstein and Tardent 1984). The high velocity of eversion and the small area of the tip of the harpoon produce extremely high pressure where the harpoon hits the prey, allowing the nematocyte to pierce even the hard exoskeleton of crustaceans (Park et al. 2017; Nüchter et al. 2006; Holstein and Tardent 1984). Studies in anthozoans have shown an analogous increase in pressure, opening of a reinforced apical structure (apical flaps), and eversion of the harpoon and tubule though this process is considerably slower in anthozoans than in medusozoans (Karabulut et al. 2022; Godknecht and Tardent 1988). In this study, we show that nematocytes from *C. americana* discharge significantly more slowly than nematocytes from *N. vectensis* but, to our surprise, ptychocytes were not significantly different from *C. americana* nematocytes. We propose the difference in firing speed may be related to the apical morphology of the capsule. A previous comparison of the capsule morphology from a variety of cnidarians demonstrated clearly that the nematocytes of tube anemones have neither an operculum nor a system of reinforced apical flaps, but rather these cells have only a thin apical cap similar to the apex seen on spirocytes and ptychocytes (Reft and Daly 2012). We propose that reinforced apical structures (opercula or flaps) allow more pressure to build up in the capsule before it is forced open, leading to a higher initial velocity of eversion. Thus, the origin of reinforced apical structures signified a major innovation in the evolution of fast firing speed in cnidocytes.

## Conclusions

Here we have shown that tube anemones have a high diversity of cnidocytes and many cell types are restricted to only certain tissues. Ptychocytes have apical sensory cones similar to those found on nematocytes, but there are no mechanosensory neuron cones in the body wall so the sensory pathway to induce firing may be different in the two tissues. Construction of the tube may involve more than just ptychocytes, as these cells are surrounded by multiciliated cells that may support the aggregation of the discharged ptychocytes into a meshwork. The morphology of the cnidocyte capsule is correlated with significant differences in speed of discharge, suggesting capsule modifications were key innovations in the diversification of cnidarians. Examining the morphologies of cells and tissues resulting from differing selection pressures for differing functions can give us a window into evolutionary pathways that give rise to diversity.

## Acknowledgements

This work made use of the Cornell Center for Materials Research shared instrumentation facility for preparation of SEM samples. Live imaging of cnidocyte discharge was performed using the Cornell Institute of Biotechnology’s BRC Imaging Facility (RRID:SCR_021741). This work was supported by the National Institutes of Health (R35GM147253-01 to LSB) and institutional funds from Cornell University. We are grateful to Gustav Paulay for his assistance with animal collection.

## Notes

### Competing Interest Statement

The authors have declared no competing interest.

